# Linked Data in Neuroscience: Applications, Benefits, and Challenges

**DOI:** 10.1101/053934

**Authors:** B Nolan Nichols, Satrajit S. Ghosh, Tibor Auer, Thomas Grabowski, Camille Maumet, David Keator, Maryann E. Martone, Kilian M. Pohl, Jean-Baptiste Poline

**Affiliations:** Center for Health Sciences, SRI International, Menlo Park, CA; Department of Psychiatry and Behavioral Sciences, Stanford University School of Medicine, Stanford, CA; McGovern Institute For Brain Research, Massachusetts Institute of Technology, Cambridge, MA; Department of Otolaryngology, Harvard Medical School, Boston, MA; MRC Cognition and Brain Sciences Unit, Cambridge, UK; Department of Radiology, University of Washington, Seattle, WA; Warwick Manufacturing Group, University of Warwick, Coventry, UK; Department of Psychiatry and Human Behavior, University of California, Irvine; University of California San Diego, USA; Helen Wills Neuroscience Institute, H. Wheeler Brain Imaging Center, University of California Berkeley

**Author notes:** co-primary authorship (B Nolan Nichols, Satrajit S. Ghosh).

## Abstract

The fundamental goal of neuroscience is to understand the nervous system at all levels of description, from molecular components to behavior. The complexity of achieving this goal in neuroscience, and biomedicine in general, poses many technical and sociological challenges. Among these are the need to organize neuroscientific data, information, and knowledge to facilitate new scientific endeavors, provide credibility and visibility of research findings, and increase the efficiency of data reuse. Linked Data is a set of principles based on Web technology that can aid this process as it organizes data as an interconnected network of information. This review examines the history, practical impact, potential, and challenges of applying Linked Data principles to neuroscience.

## Introduction

Progress in neuroscience often depends on the ability of researchers to investigate hypotheses about theories based on rigorously acquired data^1,2^. Most neuroscience research laboratories currently perform hypothesis-driven investigation using data acquired “in-house” (i.e., by the laboratory that performs the analysis). However, confining data and their analysis to a single laboratory eliminates their potential for reuse by the broader community. In recognition that neuroscience studies are not as reproducible as previously thought^3^, data reuse becomes important not only to verify results or reanalyze data, but also to aggregate data to increase sample size and boost statistical power^4^. For example, recent advances in neuroscience were only made possible by aggregating a large number of samples that are generally difficult to obtain by a single laboratory^5–7^. The difficulty of aggregating big and complex data into a form ready for analysis is a barrier to data-intensive science and a principle concern of data science. In combination with the accelerating rates of scientific data collection^8^, societal pressure for more transparency^9^, and cost effective research, the neuroscience community is now challenged to create models that increase the reuse of data for scientific discovery and to improve trust in scientific claims.

A growing number of projects, such as the Allen Institute for Brain Science^10^, the Human Connectome Project^11^, or the United Kingdom Biobank^12^ provide large datasets to the neuroscience community, and funding agencies are now often requiring data sharing^9,13^. Given the increasing size and diversity of datasets, the need to harmonize data from different sources is driving community-based efforts to describe these data in a standard way, such as the ones from the Research Data Sharing Without Barriers^14^ and the International Neuroinformatics Coordinating Facility (INCF)^15^. To improve resuse of biomedical data from the institutional level, the Office of Data Science (ODS) at the National Institutes of Health (NIH) created the Big Data to Knowledge (BD2K) initiative^16^ and the European Union started the European Life-science Infrastructure for Biological Information (ELIXIR)^17^. These initiatives foster the participation of individual laboratories or researchers in a more transparent scientific community that values Findable, Accessible, Interoperable, and Reusable (FAIR)^18^ data and research deliverables.

A key component of the FAIR principles (outlined in Box 1) is the use of machine readable metadata that provide information about data, such as the the researchers that created the dataset, the lab instruments used during data acquisition, or specific cell types contained in the data set. Scientific software could then use the machine readable metadata to track data provenance (i.e., metadata that chronicles how data were generated and processed) and foster reuse and integration of data.

### Box 1 FAIR Data Principles

The FAIR data principles^18^, developed by the FAIR working group and made available through FORCE11, proposes the use of Web standards for organizing metadata to transform how data are (re)used^25^. FORCE11 defined a set of principles that are designed to make data FAIR - Findable, Accessible, Interoperable, and Reusable^18^. Specifically, the principles are below.

To be Findable:

F1. (meta)data are assigned a globally unique and eternally persistent identifier.

F2. data are described with rich metadata.

F3. (meta)data are registered or indexed in a searchable resource.

F4. metadata specify the data identifier.

To be Accessible:

A1 (meta)data are retrievable by their identifier using a standardized communications protocol.

A1.1 the protocol is open, free, and universally implementable.

A1.2 the protocol allows for an authentication and authorization procedure, where necessary.

A2 metadata are accessible, even when the data are no longer available.

To be Interoperable:

I1. (meta)data use a formal, accessible, shared, and broadly applicable language for knowledge representation.

I2. (meta)data use vocabularies that follow FAIR principles.

I3. (meta)data include qualified references to other (meta)data.

To be Re-usable:

R1. meta(data) have a plurality of accurate and relevant attributes.

R1.1. (meta)data are released with a clear and accessible data usage license.

R1.2. (meta)data are associated with their provenance.

R1.3. (meta)data meet domain-relevant community standards.

One way of creating machine readable metadata compliant to the FAIR principle is via Linked Data^19,20^, which consists of a set of principles and technologies for interconnecting data using the infrastructure of the World Wide Web. Although 31% of all Web pages use Linked Data for tasks such as improving information retrieval^21^, adoption of the technology in neuroscience has been slow due to sociological issues, ethical concerns^22,23^, and the need for domain scientists and the data science community^24^ to agree on standards. This article reviews the 1) principles and technologies of Linked Data and how they could implement the FAIR principles, 2) existing applications of Linked Data in biomedical and neuroscience research, and 3) the challenges and limitations of applying Linked Data to scientific problems.

### What are Linked Data?

The World Wide Web was designed to be an information system for serving and linking digital documents on the Internet^26^. The Web has changed over time from a collection of static documents constructed by Web developers in the 1990’s into a Web of dynamic content driven by social media in the 2000’s (see Box 2 for details on the stages of the Web). In 2001, the Semantic Web^27^ was introduced and outlined a vision for how linking data on the Web could be used to advance sophisticated knowledge retrieval. For instance, the Semantic Web would enhance the search results for a restaurant by automatically providing links to make a reservation or display the dinner menu. In today’s world, the Semantic Web is relevant to systems like Alexa^28^, Google Now^29^, or Siri^30^. A core technology behind the Semantic Web is Linked Data, which proposes to not only use links to make a direct connection between two documents, but indicates the specific *type of relationship* that exists between these documents. These semantic links can provide information about the type of document (e.g., a scholarly article or a dataset) or other relevant metadata (e.g., the author’s name or keywords) that can be intelligently used by software applications.

#### Box 2 Evolution of the Web

Initially, the **Web (1.0)** provided a publishing platform for static documents akin to a digital version of books or newspapers for people to read and discuss. The document creators provided clickable links to other documents, much in the same way authors reference other manuscripts, but with an interactive component that enabled Web “surfing” of what are typically read-only documents.

As the **Web (2.0)** became ‘read-write,’ even novices and non-computer savvy users were able to contribute content using Web applications that make it simple to blog or comment on the work of others. In addition, pages became dynamic and started to update content without interaction from the reader.

The **Web (3.0)** is becoming a Web of Data that can be used by people and computers to access and gain knowledge from data and documents. By embedding semantic information about webpage content that computers can process, algorithms are able to automate many tasks, such as recommending content tailored to individual interests and linking related services (e.g., calendar and email). For example, a computer can follow links between Web pages, read information on the page, and display it to a user. Links within and between Web pages can contain computer processible tags or labels, hidden to the user, that provide the meaning of specific content (e.g., page title, article author, or related content). By providing links that are labeled with metadata, web documents can form a Web of Data that computer programs can use to automate tasks for users (e.g. If the web document is about a dataset, a list of appropriate analysis tools could be recommended.)

Linked Data is based on three principles with the overall goal to make it easier for people and machines to explore and publish data (Figure 1)^19^. The first principle is to use Web addresses (i.e., Uniform Resource Identifiers: URIs^31^) as identifiers, symbols, or names to refer to any concept (e.g., the Linked Data repository Wikidata^32^ represents the concept “hippocampus” using the URI: <u>http://www.wikidata.org/entity/Q48360</u>). Using URIs that start with the familiar ‘HTTP’ prefix make it possible to look up these symbols in a Web browser and retrieve additional data. The second principle requests that data is presented in a standard format so that software informed of those standards can easily process the corresponding information (see Section “Common data model” for additional information). Based on this standard, data can now be processed using a common set of tools shared among the Web community reducing the need for custom software. The third, and final, principle is that data retrieved from a URI should include links to additional data to create a *Web of Data* that eases discovery of additional information.

**Figure 1.**
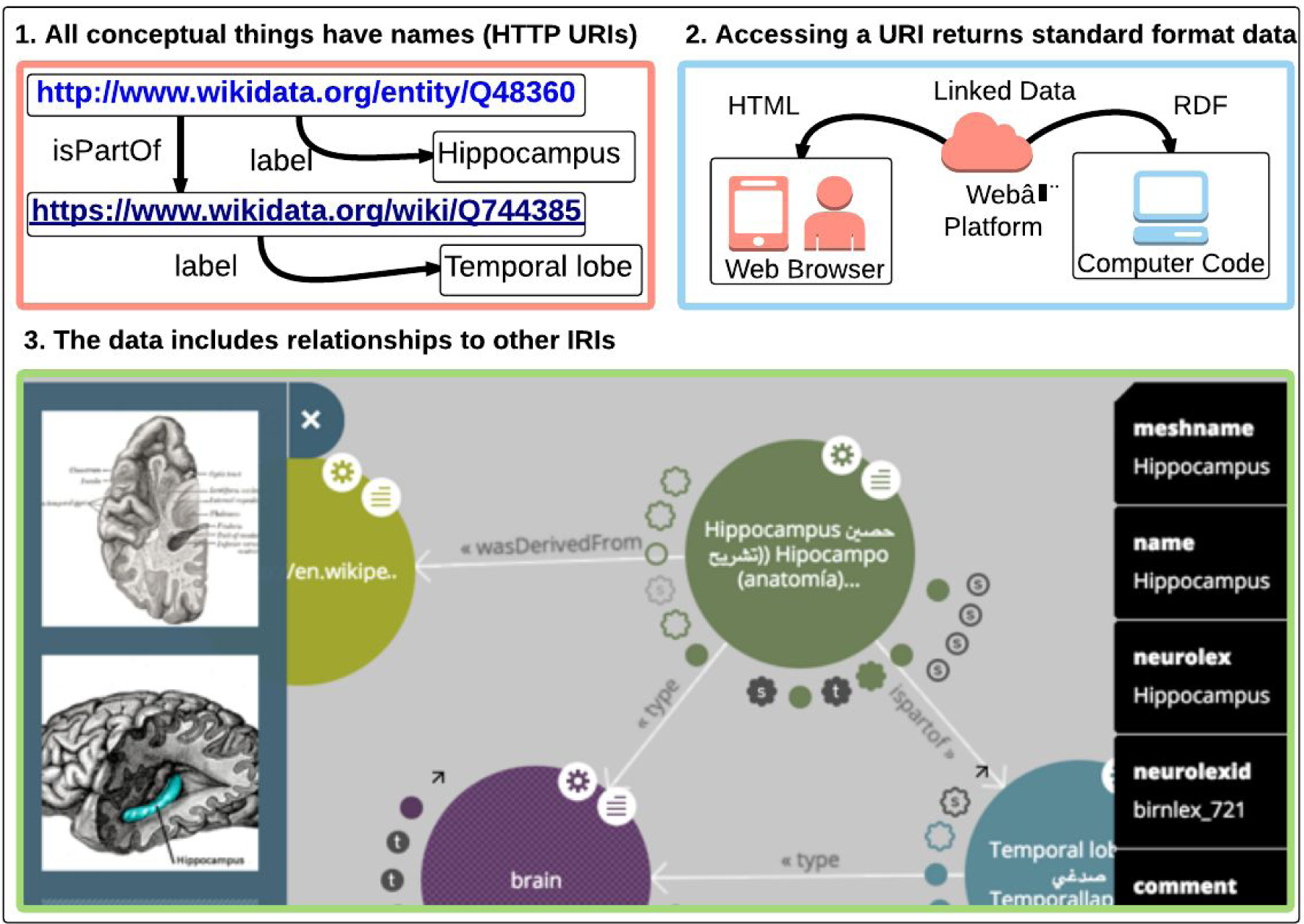
Linked Data Principles. The three Linked Data Principles^20^ allow data on the Web to be represented, found, and related. 1) Concepts should be given identifiers using HTTP URIs that 2) can be accessed using a Web browser or programming language that returns data in a standard format, such as HTML or RDF, with 3) related information encoded via links to Web accessible data. The graphical representation of the links shown in 3) was generated by the LodLive Linked Data Browser^33^. Linked Data technologies yield a powerful framework to implement the FAIR principles.

The Linked Data principles are implemented using Web technologies that are interoperable and compatible with the goals of the FAIR principles. These include: 1) a standard data model to structure data into a human and machine readable format, 2) schema languages for describing the semantics of data using standard vocabularies, 3) a query language to retrieve and filter data, and 4) security protocols to ensure data is only accessed by authorized individuals. These concepts and corresponding technologies are summarized in Figure 2.

**Figure 2.**
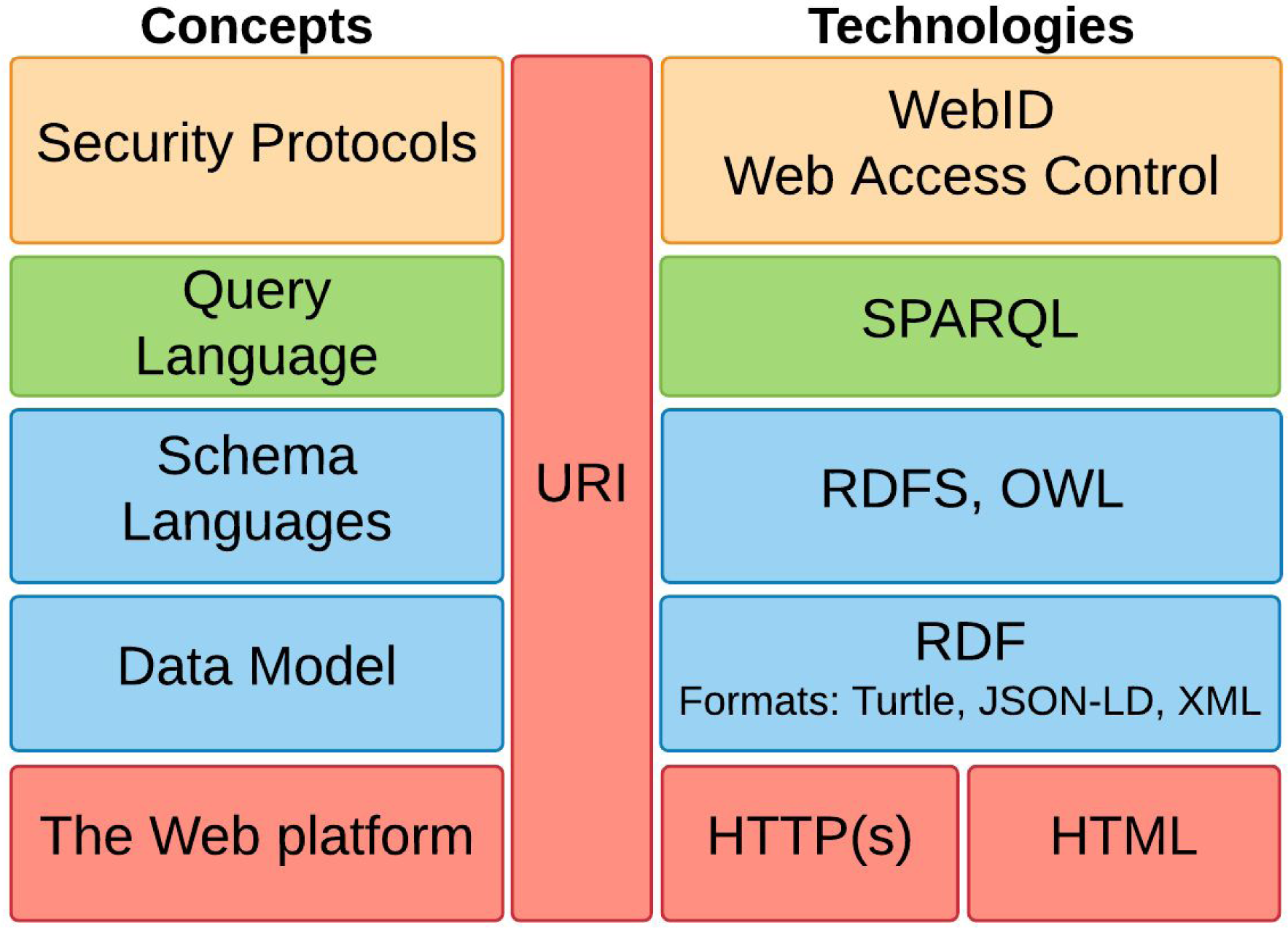
Linked Data Technologies. These Linked Data principles can be implemented using a subset of Web standards (Web platform, Data Model, Schema Languages, Query Language, and Security Protocols) to interconnect data. All listed technologies make use of Uniform Resource Identifiers (URI or equivalently URL^34^).

Applying these principles to spreadsheets, such as the one shown in Figure 3A, is an example on how Linked Data can improve the ‘readability’ of data and thus increase compliance with the FAIR principles. Spreadsheets are commonly used in neuroscience studies as they reduce the complexity of scientific data to a flat, two-dimensional data structure. They often list acronyms and derived scores, instead of the original measurements, that are known to the authors or are explicitly defined by external documents (e.g., data dictionaries). Applying the principles of Linked data can now be used to create links to other data (e.g., metadata) that fully describe the information stored in the spreadsheet. In accordance to FAIR principles, these other data needs to be easily understood by scientists, which requires the metadata to be defined according to common vocabulary designed and maintained by the scientific community. Furthermore, metadata can be used by software to automatically link to other data or metadata, which requires the use of interoperable technologies described below.

**Figure 3.**
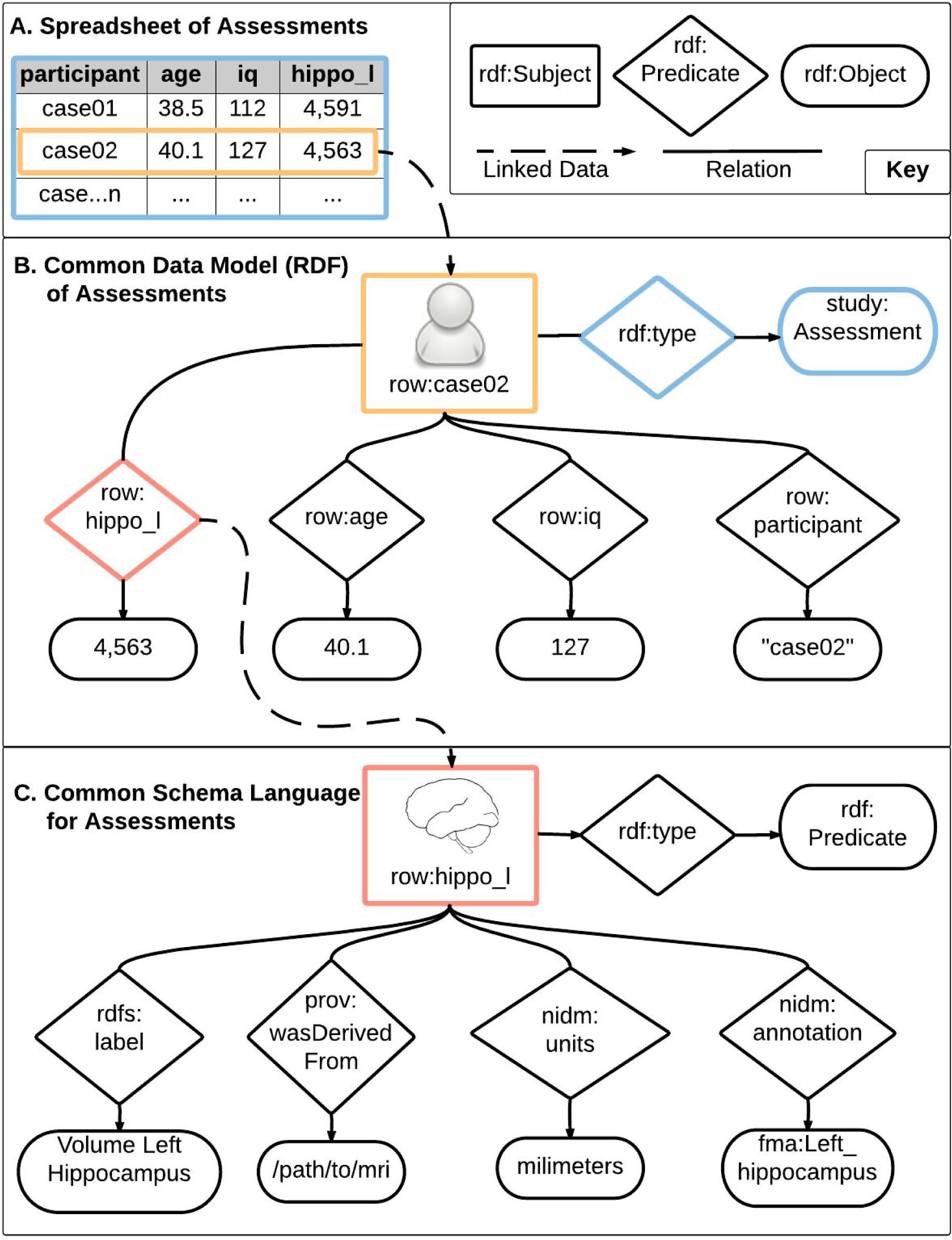
Spreadsheets on the Web. Spreadsheets are a common format for exchanging and processing tabular data in analysis packages. **A)** This spreadsheet of assessments data contains records (i.e., rows) with columns denoting a participant identifier (participant), demographic information (age), a neuropsychological test result (iq), and an anatomical measurement (hippo_l). **B)** Each cell in this tabular spreadsheet can be represented as an RDF statement (i.e., triple) with a Subject, Predicate, and Object or Value. The ‘Subject’ is denoted here using ‘row:case02’ where the ‘row:’ prefix (i.e., namespace) is shorthand for the base Web address (e.g., http://my.study.com/row#) that is expanded from ‘row:case02’ to ‘http://my.study.com/row#case02’, where one would get the information linked to this row. The columns of the spreadsheet are modeled as a ‘Predicate’ that connects the ‘Subject’ to a specific ‘Object’ or ‘Value’ to form a statement (i.e., triple). Each row of the spreadsheet is about a ‘study:Assessment’, while the remaining ‘Predicates’ (i.e.,hippo_l, age, iq, participant) connect to specific data values for each respective column (i.e., 4,563, 40.1, 127, case02). **C)** Each ‘Predicate’ is also denoted using a Web address, thus allowing it to be a node in another set of statements. These metadata can be used, for example, to decipher the term ‘hippo_l’ as “Volume Left Hippocampus”, that the measure was derived from an MRI scan, that the units of analysis were measured in millimeters, and that it is annotated with the URI ‘fma:Left_Hippocampus’ from the Foundational Model of Anatomy (FMA) ontology^35,36^. Using the RDF representation, both data and metadata can be stored in the same document or distributed across the Web.

### Standard data model

Data models define how data are connected to each other and how they are stored. A data model recommended by the W3C is the Resource Description Framework (RDF^37^), which represents information in the Web^38^ as a network of resources. RDF defines a Web-based graph database through a list of “statements” that are commonly referred to as “triples” as they are comprised of of three components: a link, also called predicate, that connects a ‘subject’ to a an ‘object’. For instance, encoding the sentence “Hippocampus is a brain region” as a RDF triple would result in “Hippocampus” being the subject, “is a” being the predicate, and “brain region” being the object. In RDF, subjects, and predicates are always encoded as Web addresses (e.g., <u>https://www.wikidata.org/wiki/Q48360</u> correspond to “Hippocampus” and <u>http://www.w3.org/2000/01/rdf-schema#subClassOf</u> to “is a”), whereas objects can be encoded as a Web address or literal values (e.g., an integer, string, etc.). With respect to the information stored in a spreadsheet, this data model can be used to augment the data with helpful metadata (e.g., the exact definition behind column names o, what data are available for each participant). Furthermore, by using Web addresses as the names for subjects, predicates, and objects, RDF datasets can link to resources on the Web that can be automatically looked up. Figure 1B provides an example of how a spreadsheet can be augmented with RDF to provide information that links to content on the Web and more closely aligns with the FAIR principles.

### Schema languages

Schema languages are used to create vocabularies that define the meaning of terms and the relationships between terms. The schema languages for Linked Data are RDF Schema (RDFS)^39^ and the Web Ontology Language (OWL)^39,40^. Analogous to the use of words to communicate concepts in natural human language, these languages use addresses on the Web (i.e., URIs) as symbols to convey meaning to data. For example, Neurolex^41^, a Linked Data application providing community curated neuroscience knowledge, uses <u>http://uri.neuinfo.org/nif/nifstd/birnlex_2092</u> as the Web address defining “Alzheimer’s disease”. These Web address-based symbols can be organized into a hierarchy that provides a formal representation (commonly referred to as a vocabulary, terminology, or ontology) of the concepts within a domain (e.g., “Neurodegenerative Disease”) and the relationships between those concepts. Creating such a vocabulary can then be used to discover related symbols. Web addresses representing “Alzheimer’s disease”, “Cerebral Palsy”, and “Parkinson disease” are examples of symbols related to “Neurodegenerative Disease”. The Web addresses representing these terms may contain structured links to specific content including definitions, synonyms (e.g., Alzheimer’s dementia), or abbreviations (e.g., AD), and included as part of a disease vocabulary (e.g., the Disease Ontology^42^ or Autism Ontology^43^)

There are already a number of resources to find specific vocabularies or annotate data with standard terms. For example, Center for Enhanced Data Annotation and Retrieval (CEDAR)^44^ is developing a user-friendly Web application to assist in annotating data with standard terms and comply with minimal information reporting guidelines. CEDAR reuses the Linked Data vocabularies hosted by BioPortal^45^. In addition, resources such as Ontobee^46^ and Linked Open Vocabularies^47^ use RDFS or OWL to describe standard terms and provide search tools. The Biosharing^48^ effort grew out of The Minimal Information for Biological and Biomedical Investigations (MIBBI)^49^, to offer an enhanced repository of digital resources that includes data standards, databases, and policies across biomedical research. The Neuroscience Information Framework has assembled a set of community ontologies of relevance to neuroscience and makes them available through web services^50^.

### Query language

Scientific data is currently stored across individual labs that cannot be queried in the same way as indexed websites (i.e., Google). By structuring scientific data using RDF and common vocabularies, searching for data could be simplified as data repositories could more easily index and archive available data. “SPARQL Protocol And RDF Query Language” was designed to answer this need. Similar to SQL, the query language for traditional databases, SPARQL can be used to select and filter one or more datasets into a form ready for analysis. The key difference is that SQL is designed for accessing tabular data while SPARQL uses graph-based pattern matching that can more flexibly access data from multiple sources that use different vocabulary. Unlike traditional databases, where schema updates are expensive operations, RDF datasets can be extended easily with new facts (i.e., triples), and SPARQL queries will continue to work.

### Security protocols

Standards for security and provenance are essential to create a balance between data availability and privacy, especially when personally identifiable information can be derived or queried using a linked Web infrastructure. Essential to creating that balance for data reuse is managing policies for data access. For example, behavioral and brain imaging data from the same participants with Autism Spectrum Disorder are distributed through multiple data repositories (e.g., NDAR^51^ and ABIDE^52^) that enforce different data use agreements. Data may also be available directly from individual participants^53–56^ or may exist on the Web without specifying a data license. For such distributed data, security protocols can protect individuals resources but are insufficient to address linkage across sources. A possible solution to enforcing policies of data shared through several resources is combining user identifiers (e.g., WebID^57^) with access control mechanisms (e.g., Web Access Control; WAC^58^). Furthermore, adding provenance records based on community-driven standards (e.g., W3C PROV^59,60^) to any published data allows tracking the data source and therefore data use constraints. Such information can be used to automatically notify data use, resource authorization, or deny access after, for example, a data use agreement expires.

To summarize, Linked Data is a Web-based architecture that consists of core components that facilitate compliance with the FAIR principles (see Box 3). Ten years after its introduction, Linked Data is becoming a well established technology on the Web with over 31% of websites having some form of Linked Data markup, which is a 22% increase since 2015^21^. For example, the Google Knowledge Graph^61^ uses Linked Data (i.e. RDFa) to optimize search performance^62^. Another example is DBPedia, a Linked Data representation of facts about people, places, and things that are displayed in information boxes on Wikipedia^63^. In addition, all metadata (e.g., titles, authors, and abstracts) published by the British Broadcasting Channel (BBC), New York Times, Nature Publishing Group (NPG), and CrossRef are made available as RDF, thus enabling computer processing of this information, and making it a part of the Linked Open Data Web. Springer Nature publishing vision embraces the use of Linked Data (<u>http://www.nature.com/ontologies</u>), and Nature Scientific Data uses ISA-Tab^64^ standard to tag publications with study description that can be represented as Linked Data and queried. Finally, a growing number of publishers are adopting the use of minimal information sets^49^, a step toward a common vocabulary and linked data principles.

#### Box 3 Linked Data and Web standards that implement FAIR

The implementation of FAIR requires metadata vocabularies and a set of Web standards. In this box, we highlight how these technologies enable data that is **F**indable, **A**ccessible, **I**nteroperable, and **R**eusable.

**Standard metadata vocabulary (RDF, OWL)**: A common data model and vocabulary allows referring to information consistently thereby enabling better search (**F**), easier integration of different datasets (**I**), and greater reuse (**R**).

**Retrieving and connecting information from the Web (URIs, HTTP)**: Uniform Resource identifiers are Web addresses. As a result the information can be viewed with a Web browser or queried automatically to improve search (**F**) but also improves access (**A**) using a common protocol (HTTP). Query across datasets (SPARQL): RDF Datasets can be distributed across the Web as files or database endpoints. SPARQL can query (**F**) and integrate information (**I** across these disparate sources.

Provenance (PROV): Each resource can be enhanced with provenance to improve specificity of search (**F**). The provenance information provides greater reusability (**R**) through improved ability to make assessments about quality.

## Applications of Linked Data in neuroscience

The use of the technologies and the adoption of the principles described above can, and to some extent already do, impact Neuroscience by providing rich data descriptions and powerful search applications as exemplified in the following.

### Leveraging bioinformatics resources

There is a growing number of Linked Data resources in the field of bioinformatics that are being effectively leveraged for neuroscience research. Resources that are essential to genomics, such as the Gene Ontology^65^ and Entrez Gene^66^ are being represented using Linked Data. Similarly, the Human Phenotype Ontology^67^ and Mouse Genome Database^68^ contain useful information for comparing across species, and have recently been integrated using Linked Data approaches. As a result, these disparate bioinformatics resources are becoming a Web of Data that can be queried as a single, integrated database and can provide answers to detailed neuroscience questions (see below).

The Bio2RDF project is an early example of integrating bioinformatics resources. Bio2RDF applies Linked Data technologies to enable queries that were not previously possible programmatically (i.e., using a query language). Using the Bio2RDF framework, Belleau et al.^69^ linked four genes to Parkinson’s disease to answer two questions in Parkinson’s disease research related to:

1. Which Gene Ontology (GO) terms describe four genes of interest (Rxr, Nurr1, Nur77, and Nor-1)?
2. Which articles mentioning these four genes of interest are related to apoptosis AND cytoplasm?

The Gene Ontology^65^ was queried for identifiers of these genes. In a second query, data collected from Entrez Gene^66^ and Pubmed^70^ sources into Bio2RDF were accessed to identify all articles with keywords and Gene Ontology annotations for “apoptosis and cytoplasm” for the genes of interest. This use case demonstrates how data from heterogeneous sources could interoperate using Linked Data, and how the results of a query could be used to get more precise and complete information on Parkinson’s disease from the literature. After identifying candidate genes, a next step may be to search across species for animal models of Parkinson’s disease. By using a query language and a specific data schema or ontology, more precise questions can be answered using existing data compared to using conventional search engines that only provide natural language documents. Another example in the Alzheimer domain is NeuroRDF, where several highly curated databases were integrated using Linked Data and then queried to identify an emerging candidate gene in Alzheimer’s disease called Macrophage Migration Inhibitory Factor^71,72^.

The Monarch Initiative^73^ is another example of the use of Linked Data in biomedical research. This computational platform uses Linked Data technologies to compare phenotypes within and across species^74^. The underlying graph database integrates and aligns cross-species gene, genotype, variant, disease, and phenotype data, such that the platform can identify animal models of human disease through similarity analysis of biochemical models. A user of this system can browse human diseases and access links to related animal models and discover related genes. For example, dystonia (<u>http://monarchinitiative.org/disease/OMIM:612067</u>) has seventeen associated phenotypes, one mouse model, one gene, and one pathway. This provides researcher with a single entry point to a network of information that previously needed to be generated with significant manual effort.

In summary, the general availability of these bioinformatics resources as Linked Data enables information to be more efficiently integrated, thus enabling better support for retrieving facts that answer biological or neuroscientific questions than manually searching across disparate databases.

### Linked Data technologies in neuroscience

While still limited, the use of the technology is also growing in neuroscience. An early example is SenseLab, which provides an entire suite of interrelated databases for molecular-level neuroscience^75^. Samwald et al. migrated these molecular data from a traditional relational database to a graph database using a semi-automated approach^75^. This process allowed data once confined to a single lab to integrate with additional resources from the community including the Brain Architecture Management System^76,77^, Gene Ontology^65^, Subcellular Anatomy Ontology^78^, and UniProt^65,79^. Harmonizing these external knowledge-bases further interconnected and enhanced the integration of information available for search^80^. Similarly, the NeuroMorpho database applies controlled terminologies to label cells and enhances the search results of queries for neuronal cell structures^81^.

Beyond single databases, aggregations of databases and datasets now use Linked Data to simplify exploration of neuroscientific data. The Neuroscience Information Framework (NIF)^81,82^ developed technology to crawl neuroscience databases^83^ and create a searchable archive of information obtained from controlled vocabularies and ontologies^81,82,84^. More recently, SciCrunch^85^ is generalizing the NIF technologies for the broader biomedical domain^85^, illustrating how these informatics solutions can reach beyond their initial domain. The National Database for Autism Research (NDAR)^51,85^ employs an ontology describing the concepts studied in autism research^43,51,85^. By incorporating the conceptual framework of an Autism ontology into NDAR, specific cognitive measures (e.g., verbal IQ) and their value ranges can be ascribed to a given concept (e.g., low verbal IQ). When rolled out over the entire resource, this enables user to browse through data conceptually, for example, by selecting all subjects labeled with ‘low verbal IQ.’ Based in China, the Linked Brain Data ^86^ is another large Linked Data project that integrates and links Web accessible brain and neuroscience data (see an example of LBD use in Box 4).

#### Box 4: Use of linked data resource example

We examined the “Linked Brain Data” (LBD, http://www.linked-neuron-data.org) resource and asked the question whether results obtained by Poldrack et al 2012 could be found using this linked data technology resource.

Poldrack et al (2012) first used a topic model to extract topics (a set of words occurring often with each others) from the neurosynth database terms, both for mental concepts and for disorders. For each of these topics, brain maps were constructed by testing at each voxel if more activity was found in papers including these terms than for other terms, constructing neural activation maps per topics. The images obtained from the mental concepts and disorders concepts were then linked using sparse canonical correlation analysis (sCCA), yielding a number of components linking mental and disorder concepts. We found that many of the disorders found by sCCA to be linked with cognitive concepts could be directly obtained using the LBD “Cognitive Functions” and “Brain diseases” relations, which are constructed with co-occurrence of terms in documents. For the first three components (canonical variates) of the sCCA, disorder terms found associated with mental constructs could be equated -or approximately equated- to LBD brain diseases in about 14 over 18 of the cases. This analysis required to match the terms used in (Poldrack 2012) and those used in the LBD, which was in general possible.

Another potential of Linked Data is the ability to retrieve information computationally that would otherwise require specific studies or significant manual effort. For example, Poldrack *et al*.^87^ used a clustering algorithm to define “topics”, also called bags of words, in cognitive functions and in disorders using the NeuroSynth database^88^ (NeuroSynth is a collection of brain imaging results spatial coordinates and associated terms from functional MRI publications). The associations between cognitive functions and disorders topics were then determined using an automated similarity analysis of the brain images associated with these topics. We found that, using the “Linked Brain Data” (LBD) resource^89^, fourteen of eighteen relations from the top three associations inferred in Poldrack et al. could be recovered, highlighting the use of a Linked Data framework as an alternative for knowledge representation and inference. See Box 4 for more details on the methods.

### Additional potential benefits of Linked Data to Neuroscience

The promises of systematically organized information date back to Otlet “réseau”^90^ in the beginning of the 20th century. While much progress has been made, his vision has not yet been realized. While the advantages of a fully implemented Web of Data are not easy to predict, and would depend greatly on the quality and extent with which data are annotated, we provide a few practical examples below of what could be achieved.

#### Aggregation, improved statistical power, and replication

With richer descriptions and linking of content that is currently disconnected, researchers could ask and obtain relevant responses to questions such as: “What manuscripts have data that supports opioid excess theories of autism?” or “What datasets are available to explore correlations between genetic and morphological variations of brain structure in autism?” (See Appendix C for a more extensive list of queries that can be enabled using richer dataset descriptors). A powerful query system would identify links to relevant data more efficiently and ease data aggregation, thus improving power of statistical analyses^4^. It would also foster replication of results on alternative datasets and integration of well documented data. The bioinformatics literature gives us an example of how Linked Data can facilitate the rapid integration of datasets, thereby reducing time spent on data management and prioritizing data analysis^69,91,92^.

#### Traceability

Linked Data can help address concerns over a lack of traceability of analyses in neuroscience. For example, Eklund et al.^93^ claims that the false positive rate in fMRI may be as high as 70% for some analyses and thus brings into question numerous brain imaging publications. This speculation could be confirmed or revoked had these manuscripts complied with the FAIR principles, by, for example, making the analysis parameters accessible to the community using Linked Data technology.

#### Acknowledgments and credits

This technology can also play a key role in crediting and acknowledging the work of others, therefore providing solutions to the, currently broken, incentive and reward system. Several publications have pointed out that: “*sharing detailed research data is associated with increased citation rate*”^94,95^. Linked Data has therefore the potential to not only help finding and documenting data but also to transform the research practices concerning the citations of research objects beyond publications and the acknowledgments of those who have provided and funded them.

## Challenges and limitations of linked data in neuroscience

In 2001, the Semantic Web^27^ intrigued researchers from information sciences to biomedicine with its vision to provide reasoning and deduction capacity to Web-based data. While its potential to solve real-world problems appeared within close reach at the time, after 15 years of research and development the vision continues to be forthcoming. Why? One issue is that many research projects tried to use complex ontology models to answer the questions of biologists and clinicians, but these projects rarely turned into useful products^96^. Furthermore, it took several years to design key Web standards (e.g., the first version of the SPARQL specification wasn’t finalized until 2008) and then develop algorithms to process graphs efficiently^97^. In summary, early attempts to realize the Semantic Web were too ambitious as they underestimated the complexity associated with comprehensive ontology modelling and the goal of deducing new facts from these models. In 2006, Linked Data took a less ambitious approach by focusing on principles for linking data (e.g., documents and metadata). Even with that focus, it took Linked Data almost another 10 years to truly take off by which time the Web industry (including Yahoo, Google, and Microsoft) realized the need for a common vocabulary to describe data in the web^21^.

While Linked data may hold many promises, the implementation faces significant technical and social challenges. The main issues are summarized below.

### Creation and maintenance of metadata vocabularies

With a field as dynamic as neuroscience, a universal agreement on a single, common vocabulary is not possible. However, the FAIR principles cannot be fully implemented without such a vocabulary or, in case of several vocabularies, their interconnection. Developing, curating, and mapping across vocabularies is at the core of the vision of Linked Data. Creating vocabularies in tandem with Linked Data faces two challenges. First, there are technical challenges related to simplifying the creation, curation, search and use of these vocabularies. Also, to ensure that a concept or an entity (e.g., “hippocampus” or “medial temporal lobe”) has a specific meaning, a unique identifier must be attributed to each concept. Community-developed vocabularies, such as those included in the Neuroscience Information Framework standard ontologies (i.e., NIFSTD)^84,98^, are an example of how to create unique identifiers for neuroscience concepts. Second, there are social challenges where individual communities must define and maintain the vocabulary of the metadata and connect it to other vocabularies. For example, several vocabularies exists for describing experimental stimuli^99–103^, but no translations exist between these vocabularies. In general, two datasets using different vocabularies cannot interoperate using common queries or analysis tools. This and other research communities using disconnected vocabularies will need to agree upon long-term strategies for stewardship over a common vocabulary or translations between vocabularies to ensure sustainability. However, there is no broadly agreed upon model for how to do this and funding agencies, publishers, communities, and institutions all have a role to play in creating linked vocabularies.

### Data processing tools will need to support Linked Data principles and provenance

In an ideal world, software processing tools would document in detail what computations have been performed, and make this information available by linking results to the original data. For instance, when a specific version of a neuroimaging tool (e.g., SPM^104^, FSL^105^, or FreeSurfer^106,107^) is used to produce derived data, information about the software, its inputs (e.g., data and parameters) and outputs needs to be recorded for reproducibility and traceability. Access to this information as Linked Data would make it possible to, for example, provide journal editors with queries that verify the analysis of a given study didn’t use software with a known error^108^ or set improper thresholds^93^. An example is the Neuroimaging Data Model (NIDM)^109^ being integrated with SPM, FSL, AFNI, and other brain imaging software tools to support exporting analysis results as a document compliant with Linked Data standards. However, given the mixture of automated and manual processing required in many neuroscience experiments, such recording is incomplete. Developing or augmenting tools to auto-document such processes will require innovative approaches to bridge the gap between legacy analysis approaches and modern workflow systems^110–113^.

### Personal information protection

Sharing human subjects data requires policies that address the potential for misuse. A common practice is to acknowledge a data use agreement that provides access to deidentified human subjects data. However, datasets that include genetic, clinical, behavioral, and imaging data can sometimes be reidentified necessitating the use of additional security measures (see Section “Security protocols”). While current Web technologies include security protocols to protect individual Web resources according to nation specific guidelines, Linked Data in combination with the Web transcending national boundaries and legal systems will require a global conversation on privacy and data reuse. In the meantime, proposed enhancements^58,114^ to the HTTP protocol are intended to support policy-aware reuse of content, including policies to protect personal information. In addition other models for data sharing are being explored. For example, the portable legal consent^115^ enables patients to effectively donate their data to science outside the guidelines of any specific study.

### Social challenges and incentives for implementing Linked Data

Social challenges around the implementation of Linked Data are arguably as important - if not more - than the technical ones, as these technologies rely on adoption by the communities. Adoption depends on factors that are both external and internal to the communities. Example of external factors comprise the recommendations or requirements from funding agencies and journal editors. For instance, NIMH now requires that data gathered using institute funds is submitted to the NDA, and NDA maintains a data vocabulary. An example of internal factors are the establishment of community guidelines on minimum reporting information^49^ such as in electrophysiology^116^, in EEG/MEG^117^, and in fMRI,^109,118–123^, or with the recent “Committee on Best Practices in Data Analysis and Sharing” of the Organization for the Human Brain Mapping, which specifies a set of concepts and information that should be documented^124^ in a publication. More generally, the social challenge is related to the current research incentive system that mostly favors innovations and new findings over the development of standards.

The adoption of Linked Data technologies will depend on software tools that provide useful services for the research community, but these tools will only be developed if there is enough Linked Data to provide useful services. Therefore Linked Data development faces a chicken and egg problem. However, the emphasis on reproducibility and traceability is increasing in biomedical research^3,125–127^, including psychology^128^, pre-clinical studies^129^, and neuroscience^4,130^. Future generations of researchers may develop and adopt a new culture where machine readable provenance information is by default included in the production of results.

## Conclusion

The rapid proliferation of neuroscience data combined with the need for a more efficient and replicable science lead to a need for standards and platforms to share, query, and access well described data. The NIH Commons^131^ project, the Open Science Framework^128^, and others are fostering community efforts that re-envision how science is conducted by prioritizing the development of a data science infrastructure aligned with FAIR. Concurrently, Web technologies following the Linked Data principles are at the core of the enhanced search and information discovery in many non scientific applications. These same technologies can provide the data science infrastructure necessary to support the principles of FAIR.

The promises of Linked Data are far from being fulfilled in neuroscience^132^. The adoption of the associated technologies faces technological and, perhaps more importantly, sociological challenges. Today, Linked Data can be adopted by teams with expertise in informatics, but not by individual researchers until user friendly tools are further developed. While tools are already emerging in neuroscience and biomedicine, with increased adoption, the Web of Linked Data could become a disruptive technology for science. It is time for neuroscience and informatics communities to use these technologies to move beyond data and information silos. Linking laboratories and research centers with scalable tools and services built on FAIR principles will not only increase access to richer and larger data sets but also yield more rapid, trusted, and findable discoveries.

## Abbreviations

**Table 1.** Acronyms

|  |  |
| --- | --- |
| AD | Alzheimer's Disease |
| BD2K | Big Data to Knowledge |
| BRAIN | Brain Research through Advancing Innovative Neurotechnologies |
| CEDAR | Center for Enhanced Data Annotation and Retrieval |
| CLARITY | Clear Lipid-exchanged Acrylamide-hybridized Rigid Imaging-compatible Tissue-hYdrogel |
| CORS | Cross-Origin Resource Sharing |
| ELIXIR | European Life-science Infrastructure for Biological Information |
| FAIR | Findable, Accessible, Interoperable, and Reusable |
| fMRI | functional Magnetic Resonance Imaging |
| FORCE11 | Future of Research Communications and e-Scholarship |
| FMA | Foundational Model of Anatomy |
| FSL | FMRIB Software Library |
| GO | Gene Ontology |
| HTML | Hyper-Text Markup Language |
| HTTP(s) | Hyper-Text Transport Protocol (secure) |
| JSON-LD | JavaScript Object Notation for Linked Data |
| LND | Linked Neuron Data |
| LOD | Linked Open Data |
| MESH | Medical Subject Headings |
| MIBBI | Minimal Information for Biological and Biomedical Investigations |
| NDAR | National Database for Autism Research |
| NIDM | Neuroimaging Data Model |
| NIF | Neuroscience Information Framework |
| NIH | National Institutes of Health |
| ODS | Office of Data Science |
| OWL | Web Ontology Language |
| PROV | W3C Specification for Provenance Information on the Web |
| SPARQL | SPARQL Protocol And RDF Query Language |
| SPM | Statistical Parametric Mapping |
| SQL | Structured Query Language |
| R2RML | Relational to RDF Markup Language |
| RDFa | Resource Description Framework in Attributes |
| RDF | Resource Description Framework |
| RDFS | Resource Description Framework Schema |
| Turtle | Terse RDF Triple Language |
| sCCA | sparse Canonical Correlation Analysis |
| SKOS | Simple Knowledge Organization System |
| SWRL | Semantic Web Rule Language |
| URL | Uniform Resource Locator |
| URI | Uniform Resource Identifier |
| W3C | World Wide Web Consortium |
| WebID | WebID Specification |
| WWW | World Wide Web |
| XML | eXtensible Markup Language |

## Acknowledgements

We thank James Brinkley, Rosemary Fama, Stephanie Sassoon, Edith Sullivan for their insightful comments on early versions of this manuscript.

SSG was partially supported by NIH grants 1R01EB020740-01A1, 3-R01-MH092380-04S2, and 1U01MH108168-01. BNN and KMP were supported in part by NIH grants AA012388, DA041123, HL127661, AA021697, the BD2K supplement AA021697-04S1 01, the Creative and Novel Ideas in HIV Research (CNIHR) Program through a supplement to the University of Alabama at Birmingham Center For AIDS Research funding (NIH P30 AI027767), which is a collaborative effort of the Office of AIDS Research, the National Institute of Allergy and Infectious Diseases, and the International AIDS Society. T.A. was supported by the Medical Research Council (United Kingdom) [MC-A060-53114]. CM has been supported by the Wellcome Trust. JBP was partially supported by NIH-NIBIB P41 EB019936, BD2K supplement AA021697-04S1, and NIH 5U24 DA039832. The Neuroscience Information Framework (MM) is supported by the NIH Blueprint via the National Institute on Drug Abuse, U24DA039832.

## Contributions

BNN, SSG, and JBP designed the overall message of the article and together with KMP developed a first draft. CM, TA, DK, MM provided input on the manuscript.

## Appendices

## Appendix A: Linked Data learning Resources and Technologies

This list provides a starting point for informaticians and database developers in neuroscience to start adopting Linked Data technologies.

Education: https://www.w3.org/standards/semanticweb/

http://www.cambridgesemantics.com/semantic-university/about-semantic-university

Libraries:

Python - <u>https://github.com/RDFLib/rdflib</u>

Javascript - <u>https://github.com/linkeddata/rdflib.js/</u>

Java - <u>http://rdf4j.org/</u>

RDF Databases:

Blazegraph (<u>https://www.blazegraph.com/</u>)

Virtuoso (<u>http://virtuoso.openlinksw.com/</u>)

Jena (<u>https://jena.apache.org/</u>)

Linked Data platforms:

Linked Data Platform: <u>https://www.w3.org/TR/ldp/</u>

Social Linked Data: <u>https://github.com/solid/solid</u>

Vocabulary search:

Ontobee (<u>http://ontobee.org</u>)

BioPortal (<u>http://bioportal.bioontology.org</u>)

NIF Vocabulary services: https://neuinfo.org/about/nifvocabularies

## Appendix B: A collection of relevant vocabularies and ontologies

NIFSTD: NIF Standard Ontology (https://neuinfo.org/about/nifvocabularies)

Gene ontology (<u>http://www.geneontology.org/</u>)

Foundational Model of Anatomy (<u>http://si.washington.edu/projects/fma</u>)

Human Phenotype Ontology (<u>http://human-phenotype-ontology.github.io/</u>)

Neuroimaging Data Model (<u>http://nidm.nidash.org/</u>)

## Appendix C: Discovery use-cases enabled by Linked Data

These use cases and queries were developed by the NIH BioCaddie^133^ project and can be answered using structured Linked Data more easily than with traditional graph databases. The Linked Data approach is event more relevant when the data are distributed across the Web.

<u>https://docs.google.com/document/d/1hVcYRleE6-dFfn7qbF9Bv1Ohs1kTF6a8OwWUvoZlDto/edit#heading=h.rr21ap8c7nwo</u>

